# A peroxide-responding sRNA evolved from a peroxidase mRNA

**DOI:** 10.1101/2021.10.18.464853

**Authors:** Madeline C. Krieger, H. Auguste Dutcher, Andrew J. Ashford, Rahul Raghavan

**Author notes:** Author for correspondence: Rahul Raghavan | |. MCK and HAD contributed equally to this work.

## Abstract

Small RNAs (sRNAs) are critical regulators of gene expression in bacteria, but we lack a clear understanding of how new sRNAs originate and get integrated into regulatory networks. A major obstacle to elucidating their evolution is the difficulty in tracing sRNAs across large phylogenetic distances. To overcome this roadblock, we investigated the prevalence of sRNAs in more than a thousand genomes across Enterobacterales, a bacterial order with a rare confluence of factors that allows robust genome-scale sRNA analyses: several well-studied organisms with fairly conserved genome structures, an established phylogeny, and substantial nucleotide diversity within a narrow evolutionary space. Using a covariance modeling-based approach, we analyzed the presence of hundreds of sRNAs and discovered that a majority of sRNAs arose recently, and uncovered protein-coding genes as a potential source for the generation of new sRNA genes. A detailed investigation of the emergence of OxyS, a peroxide-responding sRNA, demonstrated that it evolved from a 3′ end fragment of a peroxidase mRNA. Collectively, our data show that the erosion of protein-coding genes can result in the formation of new sRNAs that continue to be part of the original protein’s regulon. This novel insight provides a fresh framework for understanding how new sRNAs originate and get incorporated into preexisting regulatory networks.

**AUTHOR SUMMARY:** Small RNAs (sRNAs) are important gene regulators in bacteria, but it is unclear how new sRNAs originate and become part of regulatory networks that coordinate bacterial response to environmental stimuli. Here, we show that new sRNAs could arise from protein-coding genes and potentially be incorporated into the ancestral proteins’ regulatory networks. We illustrate this process by defining the origin of OxyS. This peroxide-responding sRNA evolved from and replaced a peroxidase gene, but continues to be part of the peroxide-response regulon. In sum, we describe the source from which OxyS, one of the most well-studied sRNAs, arose, identify protein-coding genes as a potential raw material from which new sRNAs could emerge, and suggest a novel evolutionary path through which new sRNAs could get incorporated into pre-existing regulatory networks.

## INTRODUCTION

Bacterial small RNAs (sRNAs) control gene expression by modulating translation or by altering the stability of messenger RNAs (mRNAs). sRNAs allow precise and efficient control of gene expression because they are produced quickly, regulate multiple genes simultaneously, and could degrade along with target mRNAs (1). These qualities are especially beneficial under conditions such as oxidative stress that require abrupt reprogramming of regulatory networks (2). In bacteria, oxidative stress caused by hydrogen peroxide (H_2_O_2_) is mitigated mainly by peroxidases (3, 4). For instance, a peroxidase system encoded by *ahpCF* genes is induced by the regulator OxyR when *Escherichia coli* is exposed to H_2_O_2_; OxyR simultaneously upregulates the expression of several other genes, including the sRNA OxyS that together assuage H_2_O_2_ toxicity (5–9).

OxyS is one of the most well-studied sRNAs. More than two decades of research on this sRNA has revealed many of the foundational details about sRNA-mediated gene regulation (10– 15). In *E. coli* and *Salmonella enterica*, OxyS is encoded by a gene located in the intergenic region (IGR) between *oxyR* and *argH* genes. Similar to OxyS, most sRNAs in bacteria are transcribed from genes present in IGRs; however, in recent years, numerous sRNAs that are encoded within protein-coding genes have also been identified. In such cases, sRNAs are either transcribed from promoters contained within protein-coding sequences or are generated from mRNAs by endoribonucleases that act in concert with the chaperone protein Hfq (17–22).

Despite the discovery of hundreds of sRNAs, we do not fully understand how new sRNAs originate in bacteria (23). One of the main impediments to elucidating the evolutionary histories of sRNAs is the difficulty in tracing sRNAs across large phylogenetic distances (24). Unlike proteins that are fairly easy to identify in distant bacteria, sRNAs can only be reliably detected within clusters of related microbes (25). This difficulty is due to a combination of factors, including their small size (50 to 400 nt), rapid turnover, and a lack of open reading frames (ORFs) or other features that serve as signposts (23-27). Given these constraints, an ideal group of bacteria to study sRNA evolution is the order Enterobacterales (25), which has an established phylogeny, substantial nucleotide diversity within a narrow evolutionary space, and contains well-characterized organisms with diverse lifestyles but enough similarity in genome structure to enable meaningful comparative genomics.

Here, by analyzing the prevalence of hundreds of sRNAs in more than a thousand Enterobacterales genomes, we show that most sRNAs arose recently, and that mRNAs are a potential source for the generation of new sRNAs. One sRNA that originated from an mRNA is OxyS, which evolved from a 3′ end fragment of a peroxidase mRNA. Interestingly, both the parental peroxidase and OxyS are regulated by OxyR, suggesting a novel paradigm for understanding how new sRNAs arise and are recruited into preexisting regulatory networks: Erosion of a protein-coding gene could lead to the formation of a new sRNA gene that continues to function as part of the original protein’s regulatory network.

## RESULTS

### Most sRNAs in enteric bacteria arose recently

We built covariance models for 371 sRNAs described in *E. coli* K-12 MG1655, *S. enterica* Typhimurium SL1344, and *Yersinia pseudotuberculosis* IP32953, and located their homologs across 1105 Enterobacterales genomes. The ensuing phyletic patterns of sRNA presence and absence was used to perform an evolutionary reconstruction of ancestral states using a maximum likelihood approach (**Figure 1, Table S1**). This order-wide analysis showed that 61% of sRNAs (228/371) emerged at the root of a genus or more recently (categorized as “young”). In comparison, among 148 proteins that function as gene regulators in *E. coli* and *S. enterica*, only 18% fall in this category (**Figure S1, Table S2**). The overrepresentation of recently-evolved sRNAs in our dataset indicates that sRNAs arise rapidly, probably in response to lineage-specific selection pressures. It should be noted however that the functions, if any, of most recently-emerged sRNAs have not been determined, and that nearly all sRNAs with known functions have putative origins ancestral to the root of their respective genera (belong to the “middle” and “old” categories) (**Table S1**).

**Figure 1.**
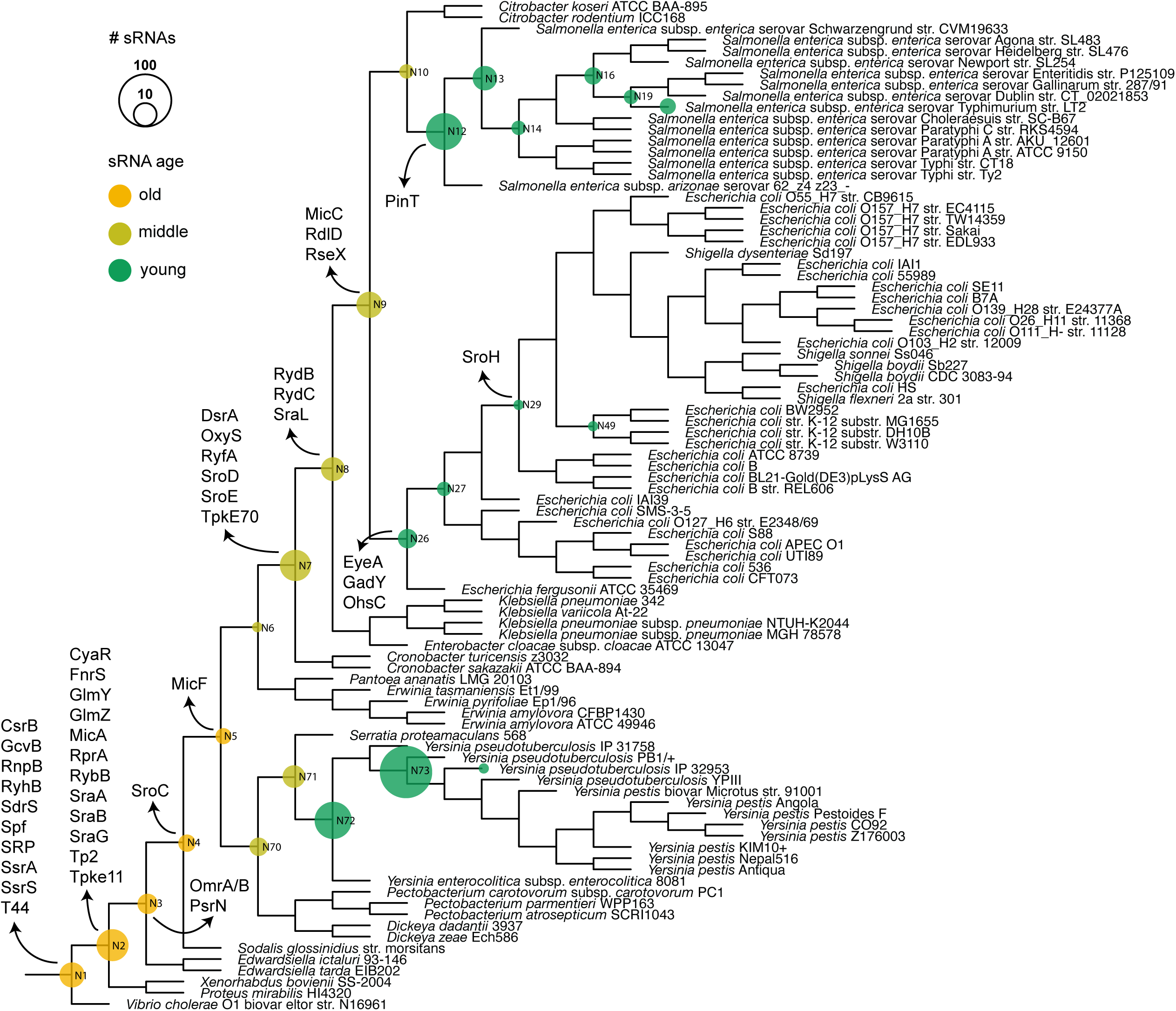
sRNA nodes of origin. sRNAs that arose at each node is depicted by circles. Size and color of each circle corresponds respectively to the number of sRNAs and their ages, as shown in the side panel. Nodes of origin of a few well-studied sRNAs are also marked.

### Protein-coding genes are potential progenitors of sRNA genes

Our covariance modeling-based search identified 62 sRNAs that were located in IGRs in the hub genomes (*E. coli* K-12 MG1655, *S. enterica* Typhimurium SL1344, or *Y. pseudotuberculosis* IP32953) but mapped to the coding strands of protein-coding genes in other Enterobacterales members (**Table S3**). A majority of the overlaps were at the 3′ ends of genes (34/62), while 18 were at 5′ ends and 10 within gene boundaries. The sRNA-ORF overlaps suggest that some of the sRNAs were originally part of mRNAs, and later evolved into independent sRNAs when the protein-coding genes decayed, leaving behind only the sRNA-encoding segments.

To better understand their evolutionary histories, we examined several sRNAs that overlapped protein-coding genes with annotated functions in more detail. This analysis revealed that OxyS, an sRNA produced in response to peroxide stress in *E. coli* and *S. enterica*, overlapped the 3′ end of a peroxidase gene in *Serratia* (**Figure 2**). The peroxidase gene is located in the same genetic context — divergent from *oxyR*, as OxyS is in *E. coli* and *S. enterica*, denoting that the sRNA likely evolved from the peroxidase gene. In addition, the promoter regions of both *oxyS* and peroxidase genes contain OxyR-binding sites, indicating that the expression of the peroxidase gene is controlled by OxyR, as shown for OxyS (6,10).

**Figure 2.**
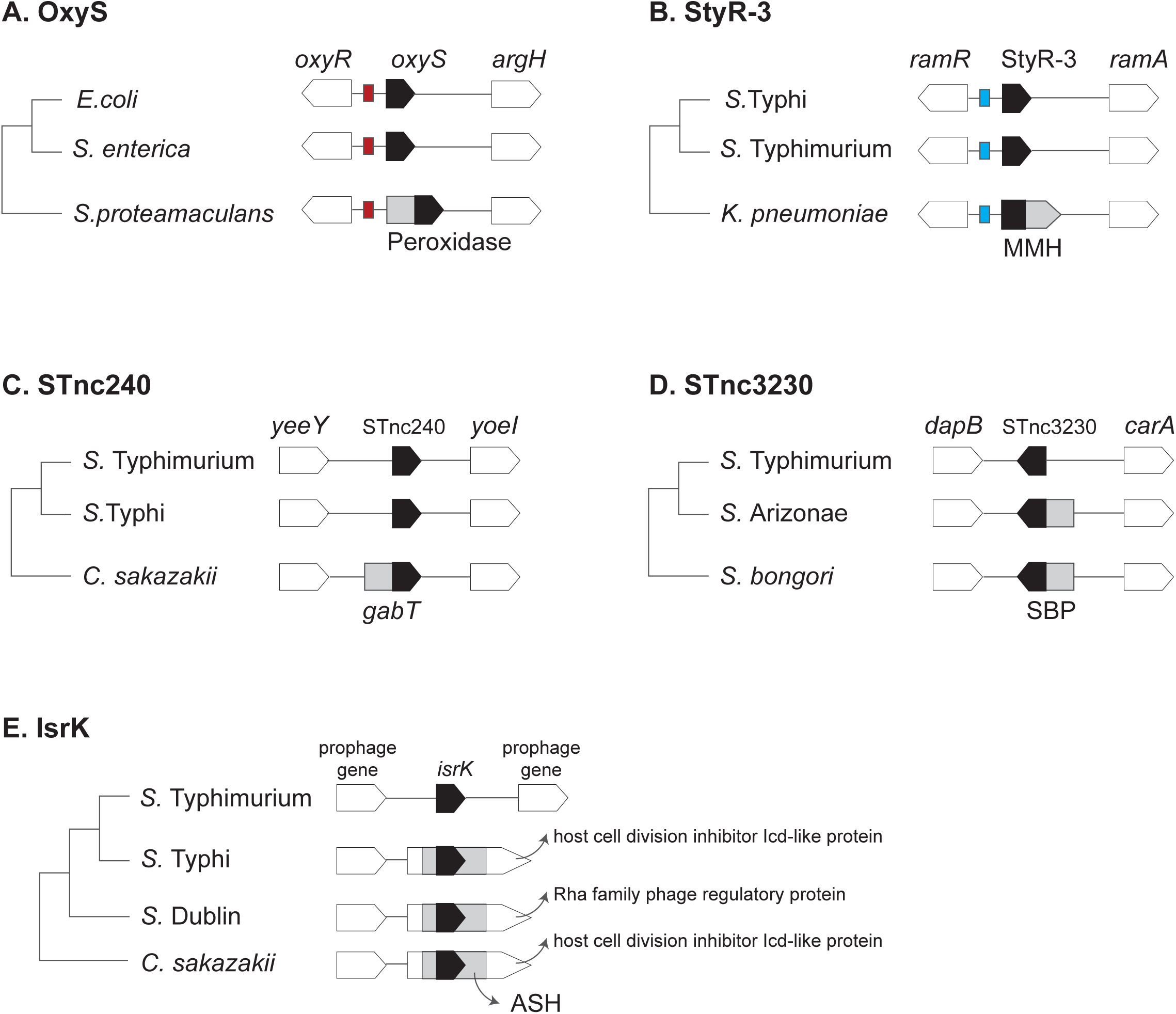
sRNAs genes replaced protein-coding genes. Several examples of sRNAs that likely originated from degraded protein-coding genes are shown. (**A**) *oxyS* genes in *Escherichia coli* and *Salmonella enterica* share sequence homology with the 3′ of a peroxidase gene in *Serratia proteamaculans*. Black arrows represent *oxyS* and its homologous sequence in peroxidase gene (grey arrow). Peroxidase and *oxyS* genes are located divergently from *oxyR*, and an OxyR-binding site (red box) is present upstream of both *oxyS* and peroxidase genes. (**B**) StyR-3 in *Salmonella enterica* Typhi and *S. enterica* Typhimurium share sequence homology with the 5′ end of an MBL-fold metallohydrolase (MMH) gene in *Klebsiella pneumoniae*. Black arrows represent StyR-3 and its homologous sequence in MMH gene (grey arrow). StyR-3 and MMH genes are located between *ramR* and *ramA* genes, and a RamR-binding site (blue box) is present upstream of both StyR-3 and MMH genes. (**C**) STnc240 in *Salmonella* species share sequence homology with the 3′ end of *gabT* gene in *Cronobacter sakazakii*, and both STnc240 and *gabT* genes are located between *yeeY* and *yoeI* genes. (**D**) STnc3230 in *S. enterica* Typhimurium and *S. enterica* Arizonae share sequence homology with the 3′ end of a sugar-binding protein (SBP) gene in *Salmonella bongori*, and both STnc3230 and SBP genes are located between *dapB* and *carA* genes. DapZ, an sRNA transcribed from within *dapB*, is not shown. (**E**) IsrK, a prophage-encoded sRNA, shares sequence homology with a region within ASH domains present in several prophage genes. Black arrows represent *isrK* and its homologous sequences in ASH domains (grey boxes).

Another sRNA that seems to be part of its parental protein’s regulatory circuit is StyR-3. This sRNA of unknown function is highly abundant in *S. enterica* (28). It shares sequence homology with the 5′ end of an MBL-fold metallohydrolase (MMH) gene in *Klebsiella*, and both StyR-3 and MMH are located divergently from the transcriptional regulator gene *ramR*. In addition, the IGR between *ramR* and StyR-3/MMH contains a RamR-binding site (**Figure 2**). The sequence similarity, homologous genetic location, and conservation of RamR-binding site suggest that StyR-3 evolved from the 5′ end of the MMH gene and continues to be under the regulatory control of the divergently encoded RamR.

A third example of an sRNA that appears to have evolved from a degraded protein-coding gene is STnc240, an sRNA with unknown function in *S. enterica*. The gene for this sRNA is located between *yoeI* and *yeeY* genes in *Salmonella* species, but in *Cronobacter*, the *yoeI*-*yeeY* IGR contains a 4-aminobutyrate-2-oxoglutarate transaminase (*gabT*) gene whose 3′ end contains a sequence that is very similar to that of STnc240 (**Figure 2**). Transcriptional regulation of *gabT* and STnc240 are not well-defined, but sequence homology and conservation of genetic location suggest that the sRNA arose from the remnants of the *gabT* gene.

An sRNA that seems to have evolved recently in *Salmonella* from a protein-coding gene is STnc3230. This “young” sRNA likely emerged from the 3′ end of a 1,3-1,4-beta-glucanase sugar-binding protein (SBP) (**Figure 2**). While both *S. bongori* and *S. enterica* Arizonae contain a gene for SBP between *dapB* and *carA* genes, STnc3230, which shares sequence similarity with 3′ end of the SBP gene, is located in this IGR in *S. enterica* Typhimurium. Interestingly, DapZ, an sRNA transcribed from the 3′ end of *dapB* gene is also located in the same IGR as STnc3230, but on the opposite DNA strand (21).

Lastly, an sRNA that seem to have evolved from within a protein-coding gene is IsrK. This prophage-encoded sRNA likely originated from the ASH domain of a bacteriophage protein-coding gene (29), and evolved to regulate the expression of a prophage-encoded anti-terminator protein AntQ (30). Similar to the origin of IsrK from a degenerated prophage gene, we have shown previously that EcsR2, an sRNA present in *E. coli*, evolved from a degraded phage tail fiber gene (26). Additionally, three sRNAs (Esr2, Esr4 and Ysr232) overlap genes that encode transposases and integrases (**Table S3**), suggesting that they arose in transposons or insertion sequences, as we showed recently for sRNAs in the pathogen *Coxiella burnetii* (31). Of these eight sRNAs, we focus on the origin of OxyS in the rest of this article.

### A peroxidase gene was replaced by *oxyS* gene in Enterobacteriaceae

In the order Enterobacterales, *oxyS* gene is present only in the family Enterobacteriaceae (e.g., *E. coli, S. enterica*), where it is located divergently from the *oxyR* gene in the *oxyR*-*argH* IGR (**Figure 3**). In contrast, a peroxidase (peroxiredoxin-glutaredoxin hybrid) gene occupies the same locus in families Erwiniaceae, Pectobacteriaceae, Yersiniaceae, Hafniaceae, and Budviciaceae.

**Figure 3.**
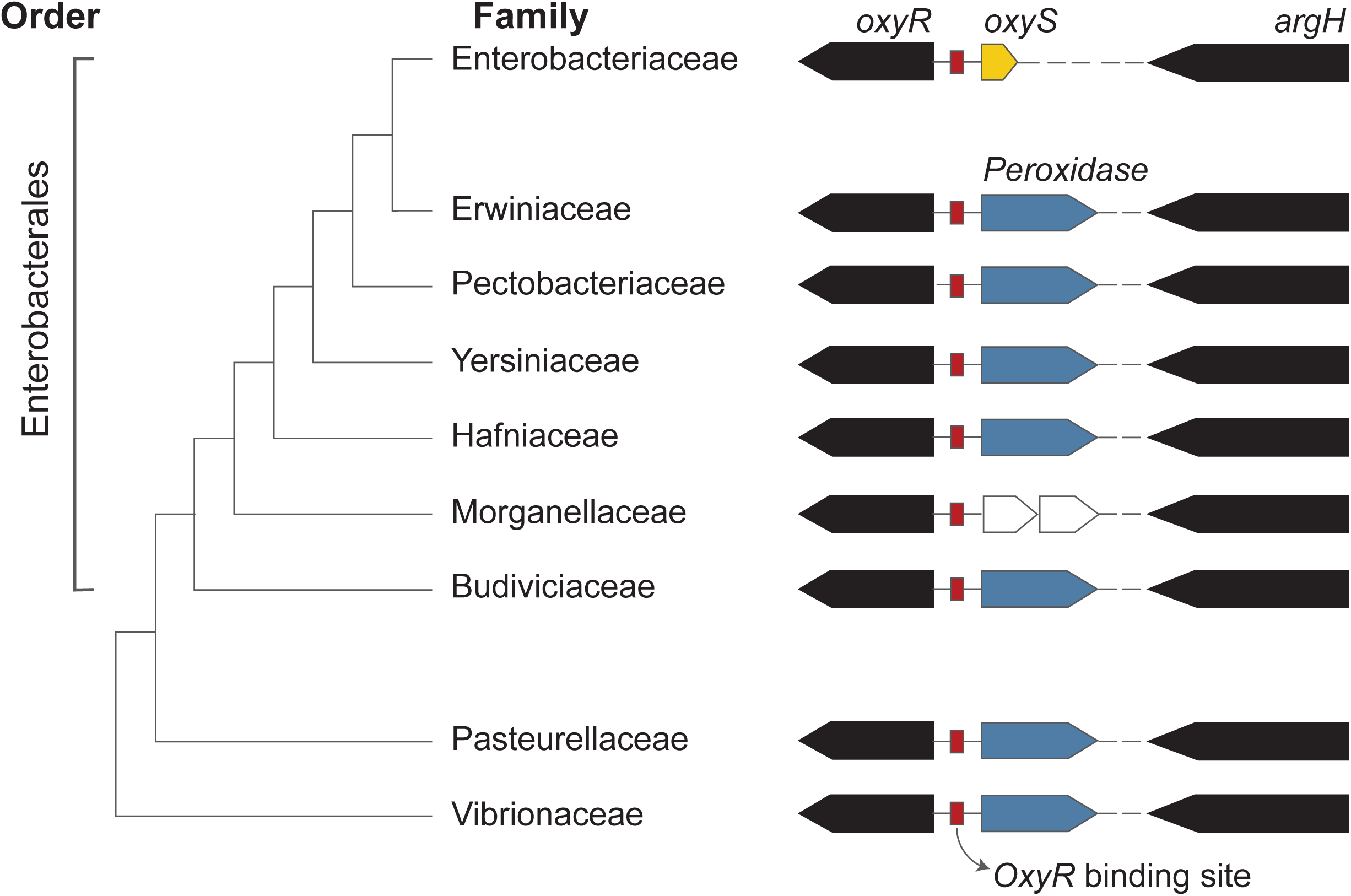
OxyS arose from a peroxidase gene. Arrangement of *oxyR*, peroxidase and *argH* genes in bacterial families within the order Enterobacterales is shown. In Enterobacteriaceae *oxyS* (yellow arrow) is found in place of the peroxidase gene (blue arrow). The intergenic region between peroxidase gene and *argH* varies between families. In members of families Pectobacteriaceae and Yersiniaceae, 3′ ends of peroxidase genes contain OxyS-like sequences (yellow arrows within blue arrows). In Morgenallaceae, transposon-associated genes (white arrows) are located in this region. Orders Pasteurellales and Vibrionales also contain the same gene arrangement as Enterobacterales members. The cladogram is based on Adeolu et al. (16).

Bacteria belonging to orders Pasteurellales and Vibrionales also contain orthologous peroxidase genes at this location (**Figure 3**). The most parsimonious explanation for this phylogenetic profile is that the peroxidase gene was present in the common ancestor of all Enterobacterales and that it was subsequently replaced by the *oxyS* gene in Enterobacteriaceae. In addition, a closer analysis of OxyS-peroxidase sequence overlap showed that the last ∼55 nt of the peroxidase coding sequence, ∼25 nt of the 3′ UTR and the intrinsic terminator likely transformed into OxyS (**Figure 4**). Based on these data, we conclude that *oxyS* gene present in Enterobacteriaceae is the remnant of the 3′ end of the ancestral peroxidase gene present in the rest of the members of the order Enterobacterales.

**Figure 4.**
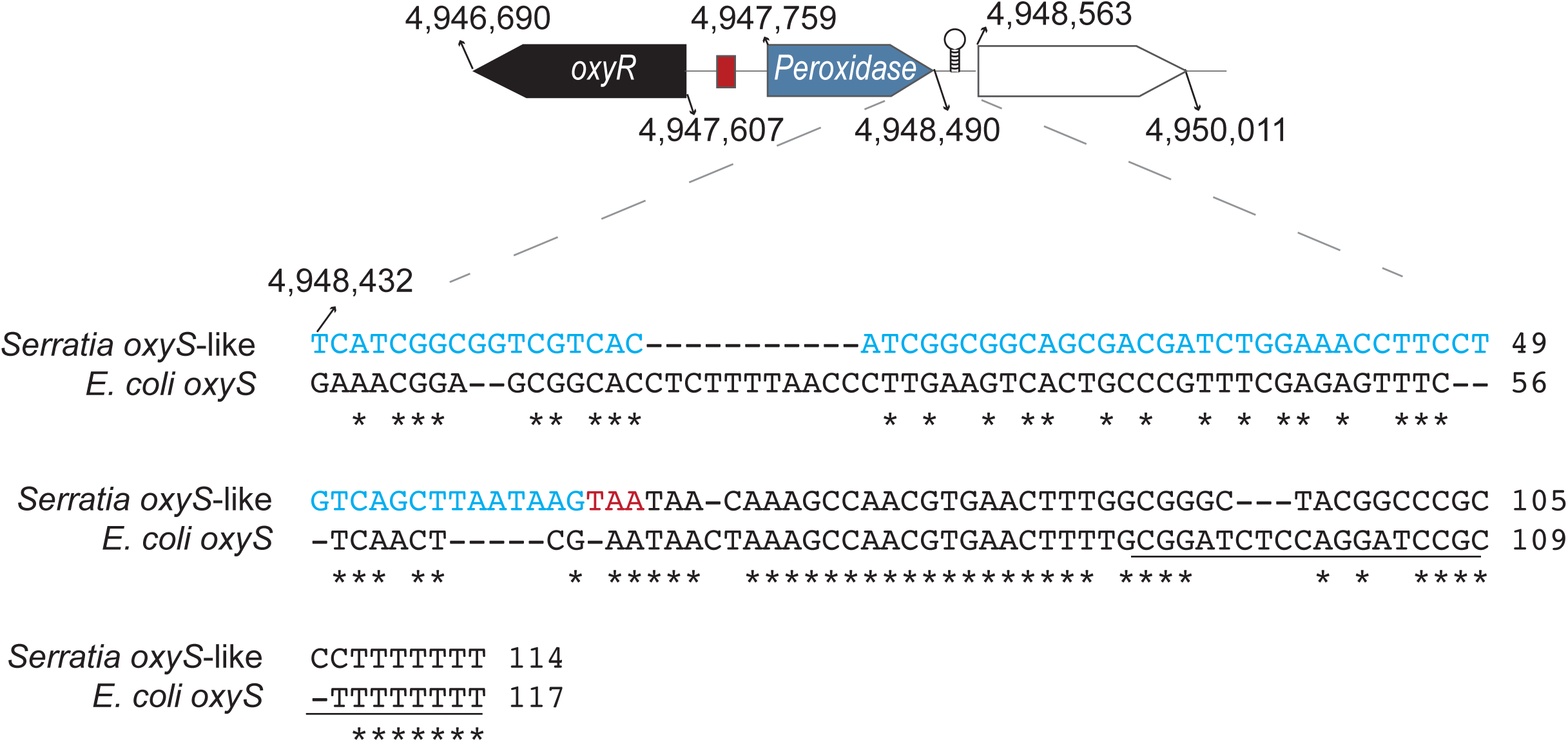
Alignment of *E. coli* OxyS with 3′ end of peroxidase gene in *Serratia*. The peroxidase gene (blue arrow) in *Serratia marcescens* ATCC13880 (CP041233) is flanked by *oxyR* (black arrow) and a dihydrolipoyl dehydrogenase gene (white arrow). ClustalW alignment of *oxyS* gene in *E. coli* MG1655 (NC_000913.3) with 3′ end of peroxidase gene is shown at the bottom. Nucleotides in blue are part of the peroxidase coding sequence, the stop codon is in red, and the predicted Rho-independent terminator sequence is underlined. The intergenic region between peroxidase and *oxyR* genes contains putative OxyR-binding sites (red square).

### Exposure to H_2_O_2_ induced peroxidase expression and production of mRNA fragments

Similar to *oxyS*, the peroxidase gene is located divergently from *oxyR* and the IGR between the two genes contain putative OxyR-binding sites (**Figure 3, Figure S2**). To test whether the expression of the peroxidase gene is induced by H_2_O_2_, we exposed *Serratia marcescens* (family Yersiniaceae), *Edwardsiella hoshinae* (family Hafniaceae), and a representative from outside of Enterobacterales: *Vibrio harveyi* (order Vibrionales, family Vibrionaceae) to 1mM of H_2_O_2_ for 10 minutes. As observed for OxyS in *E. coli* (10), exposure to H_2_O_2_ significantly induced the expression of the peroxidase gene in all three bacteria (**Figure S3**), indicating that it is also regulated by OxyR. Additionally, peroxidase mRNAs in all three bacteria produced small 3′ fragments that correspond to the region from where OxyS emerged (**Figure 5**). Although the size of the cleavage products are not identical in the three bacteria, the fragmentation process itself appears to be a trait ancestral to all Enterobacterales, thus providing the raw material from which OxyS eventually evolved in an Enterobacteriaceae ancestor.

**Figure 5.**
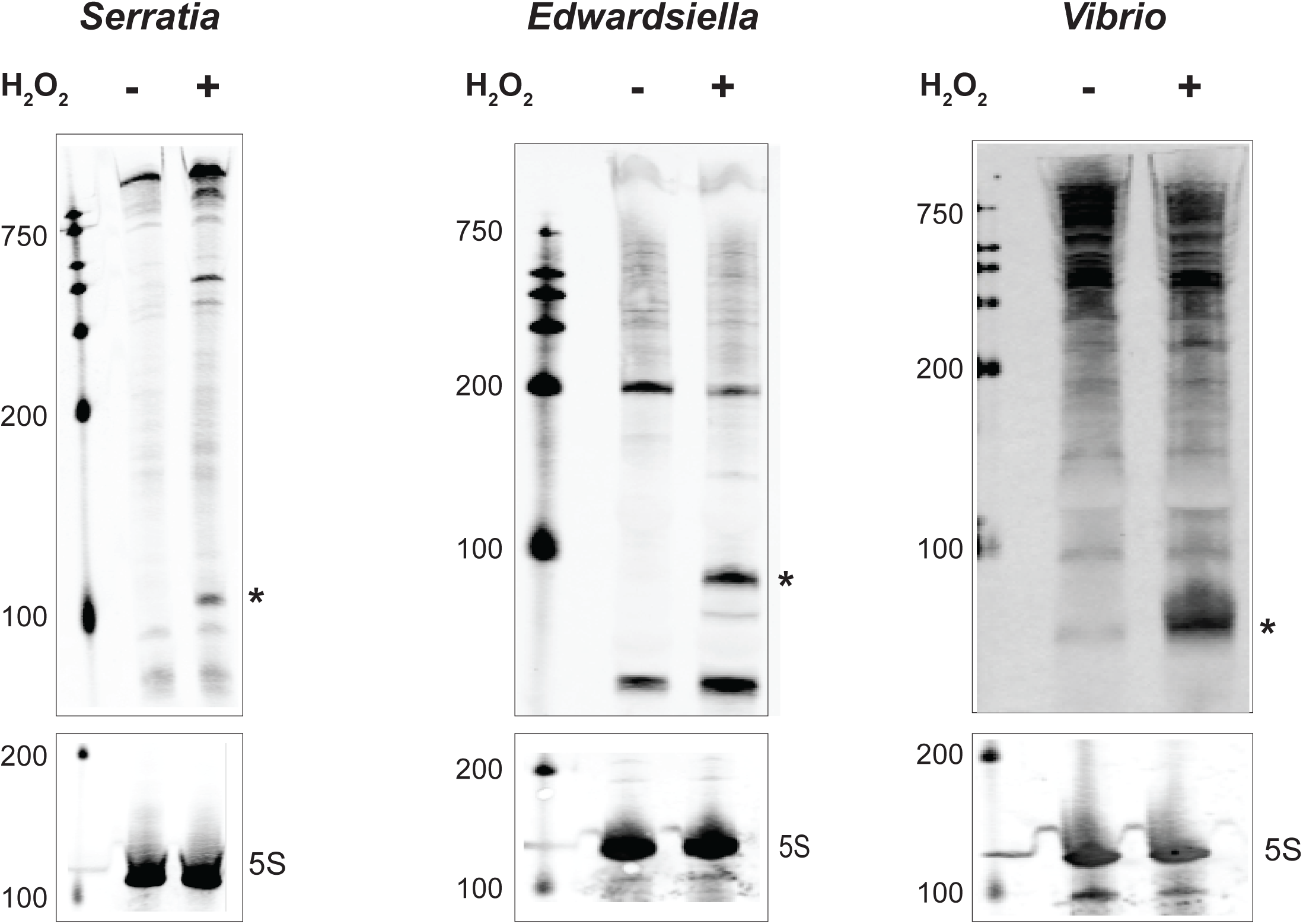
Peroxidase mRNA fragmentation. Short 3’ end fragments (marked with *) were cleaved from peroxidase mRNAs in *S. marcescens, E. hoshinae*, and *V. harveyi*. “+” indicates samples exposed to 1mM of H_2_O_2_ for 10 minutes, and non-exposure controls are shown with “-”. Northern blotting was performed with probes that bind to the 3′ ends of peroxidase mRNAs.

## DISCUSSION

Despite their importance to bacterial physiology and virulence, the evolutionary processes that produce new sRNAs are not well understood. One of the main reasons for this lack of clarity about sRNA origination is that unlike protein-coding genes, sRNA genes are difficult to trace across large phylogenetic distances (24). Following up on previous research that showed that enteric bacteria are at optimum distances from one another to effectively investigate sRNA prevalence (25), we traced the presence of hundreds of sRNAs across Enterobacterales and show that a majority emerged recently in a lineage-specific manner. This observation fits with earlier findings that showed that sRNAs evolve rapidly in bacteria and are typically genus- or species-specific (26, 32-35). Interestingly, we found that most well-studied sRNAs belong to “middle” and “old” age groups. A similar observation was made by a study that examined the evolutionary histories of 58 experimentally validated sRNAs in *E. coli* (36). Although specific age categories were not assigned in that study, when we classified the sRNAs into three age groups based on the distance of gain-node from *E. coli* (old: >0.1; middle: 0.001 - 0.009; young: 0.0001 - 0.0009), 51/58 sRNAs were deemed to be old or middle-aged (**Table S4**). This biased representation is probably due to the propensity of older sRNAs to be expressed at high levels, thereby making them more amenable to discovery and experimental validation (26, 37).

We have previously shown that new sRNAs arise *de novo* and from degraded bacteriophage- and transposon-associated genes (23, 26, 31, 33). In this study, we report that protein-coding genes could serve as a raw material for sRNA biogenesis and support this conclusion by showing that OxyS, a peroxide-responding sRNA, originated from a peroxidase gene. OxyS was first noticed by researchers because it is transcribed divergently from the *oxyR* gene that encodes a transcriptional regulator that orchestrates *E. coli*’s antioxidant response (10). The IGR between *oxyS* and *oxyR* genes contains OxyR-binding sites from where the oxidized form of OxyR induces the expression of OxyS in response to H_2_O_2_ exposure (10,38-40). We show that the *oxyR-oxyS* gene arrangement is present only in the family Enterobacteriaceae, whereas a peroxidase gene, whose expression is also induced by H_2_O_2_, occupies the genetic locus next to *oxyR* in other members of the order Enterobacterales. The evolutionary process that led to the replacement of the ancestral peroxidase gene by *oxyS* gene could have occurred through two possible routes (**Figure 6**). In one, the mRNA 3′ end fragment gained a regulatory function, which resulted in its retention when the rest of the peroxidase gene was pseudogenized and later deleted in Enterobacteriaceae. Alternatively, the peroxidase gene was pseudogenized first, but continued to be transcribed producing the 3′ end fragment, which gained a regulatory function and was retained when the rest of the gene was deleted.

**Figure 6.**
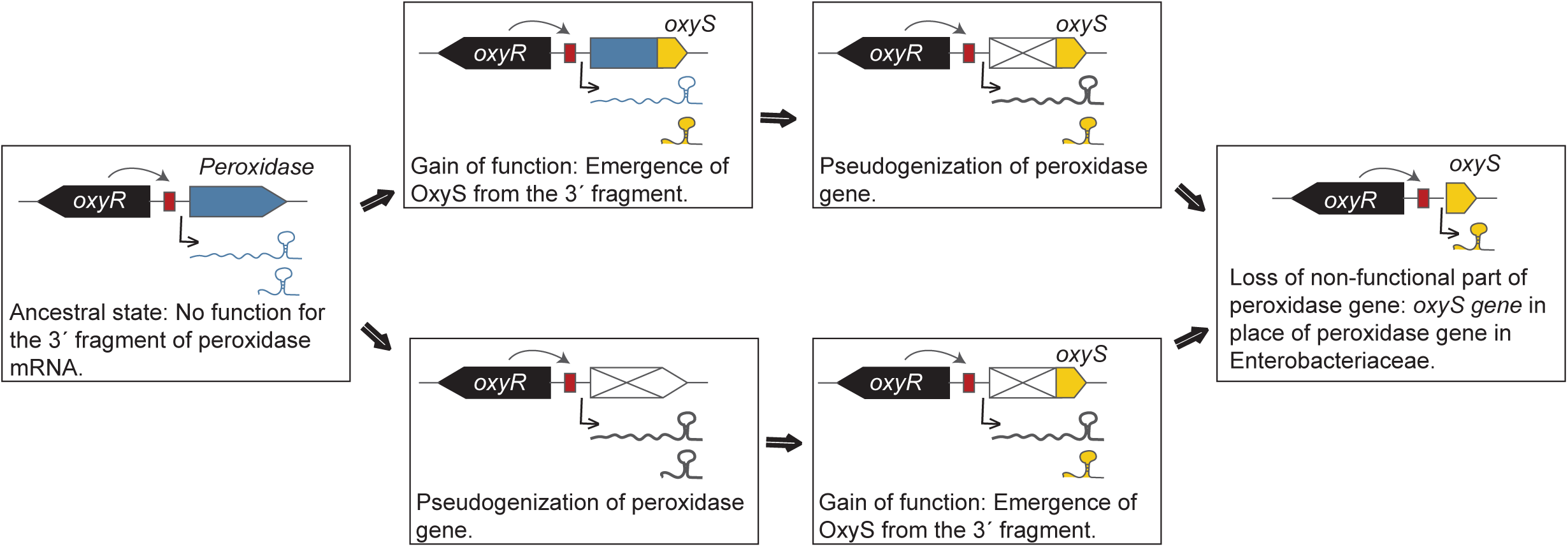
Two possible routes of OxyS evolution. The 3′ end fragment produced from the ancestral peroxidase mRNA was likely non-functional. Top path: In an Enterobacteriaceae ancestor, functional OxyS emerged as a 3′ end-derived sRNA prior to the pseudogenization of the peroxidase gene. Bottom path: OxyS emerged at the 3′ end of an RNA transcribed from a pseudogenized peroxidase gene. Ultimately, the non-functional part of the peroxidase gene was deleted from the genome, resulting in the formation of the *oxyS* gene. OxyR binds to sites (red boxes) located in the IGR and regulates the expression of peroxidase and *oxyS* genes.

An unresolved question in the field of sRNA biology is how new sRNAs become incorporated into regulatory networks. This study provides a possible explanation: An sRNA arising from a protein-coding gene would inherently be part of the parental protein’s regulon. For instance, in either scenario in **Figure 6**, the mRNA fragment that gave rise to OxyS would have been produced as part of the OxyR regulon even before it gained any function. Thus, when functional OxyS later emerged from that fragment, it was already part of the OxyR regulatory network. Similar to the origin of OxyS, 3′ ends of mRNAs appear to be the most favorable location for sRNA genesis (34/62 in our dataset) probably due to the presence of intrinsic terminators that improve RNA stability and promote Hfq binding, two factors that are critical to sRNA evolution and function (27, 41-44). The next best seem to be 5′ ends (18/62), likely due to proximity to promoter regions that regulate transcription, with middle region being the least likely (10/62) to contribute to sRNA evolution. Irrespective of this apparent difference, any part of an mRNA could potentially evolve into a regulatory RNA, as shown in a recent study that demonstrated the generation of sRNA-like transcripts from 5′, middle and 3′ segments of mRNAs in *E. coli* (20). In sum, because bacterial genomes contain numerous protein-coding genes in various stages of decay (45, 46), they could function as a rich resource from which new sRNAs could arise rapidly in response to lineage-specific environmental pressures.

## MATERIALS AND METHODS

### Determining sRNA presence across the order Enterobacterales

A list of candidate-sRNAs in *E. coli* K-12 MG1655 (NC_000913.3), *S. enterica* Typhimurium SL1344 (NC_016810.1), and *Yersinia pseudotuberculosis* IP32953 (NC_006155.1) were compiled from previously published studies (37, 40, 47). Several exclusion criteria were used to remove spurious and redundant sRNAs: (i) sRNAs under 60 nucleotides in length, (ii) sRNAs that overlapped each other by more than 10 nucleotides, (iii) sRNAs that were present in multiple copies, (iv) RNAs that were identified as cis-acting in the Rfam database (48), and (v) sRNAs that shared 95% or more nucleotide identity over at least 60 nucleotides of their lengths. For each sRNA of interest, a representative sequence from each hub genome was used as the query for a BLASTn (wordsize 7, maxdbsize 100Kb, dbsize normalized to 4Mb, evalue ≤1e-5) using BLAST v2.7.1 against a database of 1105 Enterobacterales genomes (**Table S5; Figure S4**) that met the following criteria: (i) full genome sequence was available on GenBank, and (ii) the genome was within 0.08 16S rDNA pairwise distance from the hub species (**Table S6**). Drawing on guidance from previous studies (24, 25), hits with pident >65% covering at least 95% of the length of the original query served as seed sequences from which to construct a covariance model. Candidate hits were next binned by percent identity, and a randomly selected set of sequences (one from each percent identity bin) were chosen to serve as a seed sequence for the covariance model. These sequences were aligned using ClustalW, and the Infernal suite of tools (v1.1.2) was used for subsequent covariance model construction (cmbuild), calibration (cmcalibrate), and homolog searches (cmsearch) (49). Models were constructed from the BLAST-derived seed sequences using cmbuild, while cmscan was used to identify sRNAs already represented by existing Rfam models. These newly constructed models, plus the existing Rfam models, were then used in parallel to search the 1105-genome database for homologs. For sRNAs that were represented by an existing Rfam model, cmsearch results from this model were compared to that from the newly constructed model, and the model that yielded more hits was selected for continued iteration. Results from cmsearch with an e-value <1e-5 were used to add unrepresented sequences to the query model, which was then refined, recalibrated, and used for another round of cmsearch. This process was repeated for each sRNA until a cmsearch with its corresponding model failed to yield new unrepresented sequences. In order to ensure that any two models were not yielding the same set of hits, results from cmsearch with the finalized models were compared across sRNAs; models with redundant hits were omitted, as were any models that yielded >1e4 hits. An sRNA gene was considered present in a given organism if a hit of e-value <1e-5 was found on its chromosome and/or plasmid. All resultant hits were cross-checked by genome location to ensure that a given hit was not represented more than once in the final results. Presence/absence data for all 1105 organisms was collected, but only data for 89 Enterobacteriaceae, plus *Vibrio cholerae* El Tor str. N16961 as an outgroup, was used for downstream phylogenetic analyses.

### Evolutionary reconstruction

Enterobacterales phylogenetic tree was downloaded from MicrobesOnLine, and node of origin for each sRNA was determined using the Gain and Loss Mapping Engine (GLOOME) as described previously (36, 50). For sRNAs present in a single hub genome (*E. coli* K-12 MG1655, *S. enterica* Typhimurium SL1344 or *Y. pseudotuberculosis* IP32953), the determined gain node was the most ancestral node with a posterior probability of ≥0.6, where all nodes leading from this ancestor to the hub genome had a posterior probability ≥0.6. If an sRNA was present in more than one hub genome, and the most recent last common ancestor (LCA) of the hub genomes in which it was present had a posterior probability ≥0.6, the gain node was the most ancestral node with a posterior probability ≥0.6, where all nodes leading from this ancestor to the aforementioned LCA had a posterior probability ≥0.6. sRNAs that emerged at the root of a genus or more recently were classified as “young” (n=228), those present at the last common ancestor of all three hub genomes were deemed to be “old” (n=57), and sRNAs that emerged in between the two age groups were considered “middle” (n=73). If the last common ancestor of the hub genomes in which the sRNA was present did not have a posterior probability of ≥0.6, ages of these sRNAs were considered undetermined (n=13).

### Regulatory protein tree

Using the QuickGO annotation table (51), GO:0010629 (negative regulation of gene expression) and its child terms were utilized to identify regulatory proteins in *S. enterica* Typhimurium LT2 and *Escherichia coli* K-12 MG1655. Protein sequences (UP00000104, UP000000625) were downloaded from Uniport (52). Redundancies among the regulatory proteins from the two different species were identified using a BLASTp of the regulatory protein sequence set against itself (pident >80, e-value <1e-10). Homologs of this final set of query proteins were identified using BLASTp (e-value <1e-10) against a database of protein sequences from the genomes used for phylogenetic analysis, yielding a presence/absence matrix. Node of origin was determined using GLOOME as described above.

### Finding *oxyR*-*argH* IGRs

To find *oxyR* and *argH* orthologs, tBLASTn searches were carried out with OxyR and ArgH sequences from *E. coli* K-12 MG1655 against Enterobacterales genomes (e-value ≤10e-10, percent positive ≥60%, percentage alignment length ≥60%). We corroborated the tBLASTn hits by confirming that they contain Pfam domains PF03466.20 and PF00126.27 (for OxyR), and PF00206.20 and PF14698.6 (for ArgH) (53). We then compiled a list of bacteria that have both *oxyR* and *argH* genes and obtained the nucleotide sequences between the two genes using the Entrez E-utilities tool.

### Identifying 5′ neighbors of *oxyR* and OxyR-binding sites

Using BioPython (54), we first determined the direction of the gene next to *oxyR*’s 5′ end. If the neighboring gene was oriented divergently, we determined the identity of the encoded protein using Pfam as described above. All Enterobacterales and Vibrionales (except Morganellaceae) included in this study contained a peroxidase gene with Glutaredoxin (PF00462) and Redoxin (PF08534) domains in this locus. To identify putative OxyR-binding sites located in the IGR between *oxyR* and its neighbor, we extracted 50 bp at the 5′ end of *oxyR* along with 100 bp of the adjoining IGR from each bacterium. Sequences were aligned with MUSCLE (55) and the 50 bp *oxyR* sequence was trimmed from each sequence. Using Multiple Em for Motif Elicitation (MEME) (56), we first detected ∼37 bp palindromic sequences in IGRs, and then used the output from MEME in Find Individual Motif Occurrences (FIMO) (57) to identify putative OxyR-binding site in each bacterium. Sequence logos were generated using WebLogo 3 (58).

### Bacterial growth and gene expression

Bacteria were inoculated 1:100 from an overnight culture into fresh media. *S. marcescens* ATCC 13880 was grown at 37°C in Lysogeny broth (LB), *E. hoshinae* ATCC 35051 at 26°C in LB, and *V. harveyi* ATCC 43516 at 26°C in Marine Broth, all shaking at 200 rpm. All cultures were grown to an OD600 value of 0.4-0.5, and split into two. One half was allowed to grow under the same conditions for 10 minutes, while the other half was exposed to 1mM of H_2_O_2_ for 10 minutes. RNA Stop Solution (5% phenol, 95% ethanol) was added and total RNA was extracted using TRI reagent (Thermo Fisher Scientific). RNA was treated with TURBO DNase (Thermo Fischer Scientific), and was either sent to Yale Center for Genome analysis for RNA sequencing (RNA-seq), or cDNA was synthesized, and quantitative PCR (qRT-PCR) was performed. PCR primers used in this study are listed in **Table S7**. RNA-seq reads were processed as described previously (59), and deposited in NCBI (PRJNA665492).

### Northern blot

RNA samples were loaded onto either 6% or 10% TBE-Urea Gel (Thermo Fischer Scientific) with a biotinylated RNA ladder (Kerafast). Gels were run in 1x TBE buffer at 180V for 60 minutes (6% gels) or 180V for 80 minutes (10% gels). Membranes and filter paper were pre-soaked and RNA was transferred to a Biodyne B Nylon Membrane (Thermo Fischer Scientific) overnight. Membranes were UV-crosslinked using a Stratalinker 2400 UV Crosslinker (1200 mj) and RNA probes (**Table S7**) were hybridized overnight at 45°C with rotation. Hybridization solution was removed and membranes were washed, blocked in Licor Intercept Blocking Buffer and treated with Streptavidin-IRDye 800 CW and examined on a Licor Odyssey scanner.

## ACKNOWLEDGEMENTS

We thank Samantha Fancher for assistance with *S. marcescens* experiments, and Jim Archuleta for bioinformatics support. We are grateful to Boris Görke, Svetlana Durica, Katarzyna Bandyra and Ben Luisi for providing us with RNase E.

## FUNDING INFORMATION

This project was supported in part by NIH grants AI133023 and DE028409 to R.R.

## CONFLICT OF INTEREST

The authors have no conflicts of interest to declare.

## Notes

### Competing Interest Statement

The authors have declared no competing interest.

