## Supplemental figures for "A peroxide-responding sRNA evolved from a peroxidase mRNA"

SUPPLEMENTAL FIGURE LEGENDS

**Figure S1. Regulatory proteins in enteric bacteria are older than sRNAs.** Regulatory proteins that arose at each node is depicted by circles. Size and color of each circle corresponds respectively to the number of proteins and their ages, as shown in side panel.

**Figure S2. Predicted OxyR binding sites in Enterobacterales members**. Predicted OxyR-binding sites located between *oxyR* and peroxidase/*oxyS* genes in one representative taxa per family.

**Figure S3. $H_2O_2$ induced peroxidase expression in *S. marcescens*, *E. hoshinae*, and *V. harveyi*.** Top panels: RNA-seq reads that mapped to peroxide genes (orange arrows) in *S. marcescens*, *E. hoshinae*, and *V. harveyi* when exposed to 1mM of $H_2O_2$ for 10 minutes. Middle panels: RNA-seq data from control (no $H_2O_2$ exposure) samples. Bottom panels: Confirmation of peroxidase induction by $H_2O_2$ via RNA-seq (*S. marcescens*) or qRT-PCR (*E. hoshinae*, and *V. harveyi*).

**Figure S4. Taxon representation within the 1105-genome search space.** Phylogenetic tree showing representation of the twelve most abundant genera among the 1105 genomes used to construct covariance models. Size of circle denotes number of genomes per genus, as shown in the side panel. 16S pairwise distances between representative species from each genus can be found in Table S6.

#### SUPPLEMENTAL TABLES

**Table S1.** Predicted age of sRNAs.

**Table S2.** Predicted age of regulatory proteins in *E. coli* and *S. enterica*.

**Table S3.** sRNAs that overlap ORFs.

**Table S4.** Age estimation of sRNAs analyzed in Peer and Margalit 2014 (36).

**Table S5.** NCBI accession numbers for Enterobacterales genomes used in this study.

**Table S6.** 16S pairwise nucleotide distances between twelve representative species spanning the 1105-genome search space.

**Table S7.** PCR primers used in this study.

### Figure S1

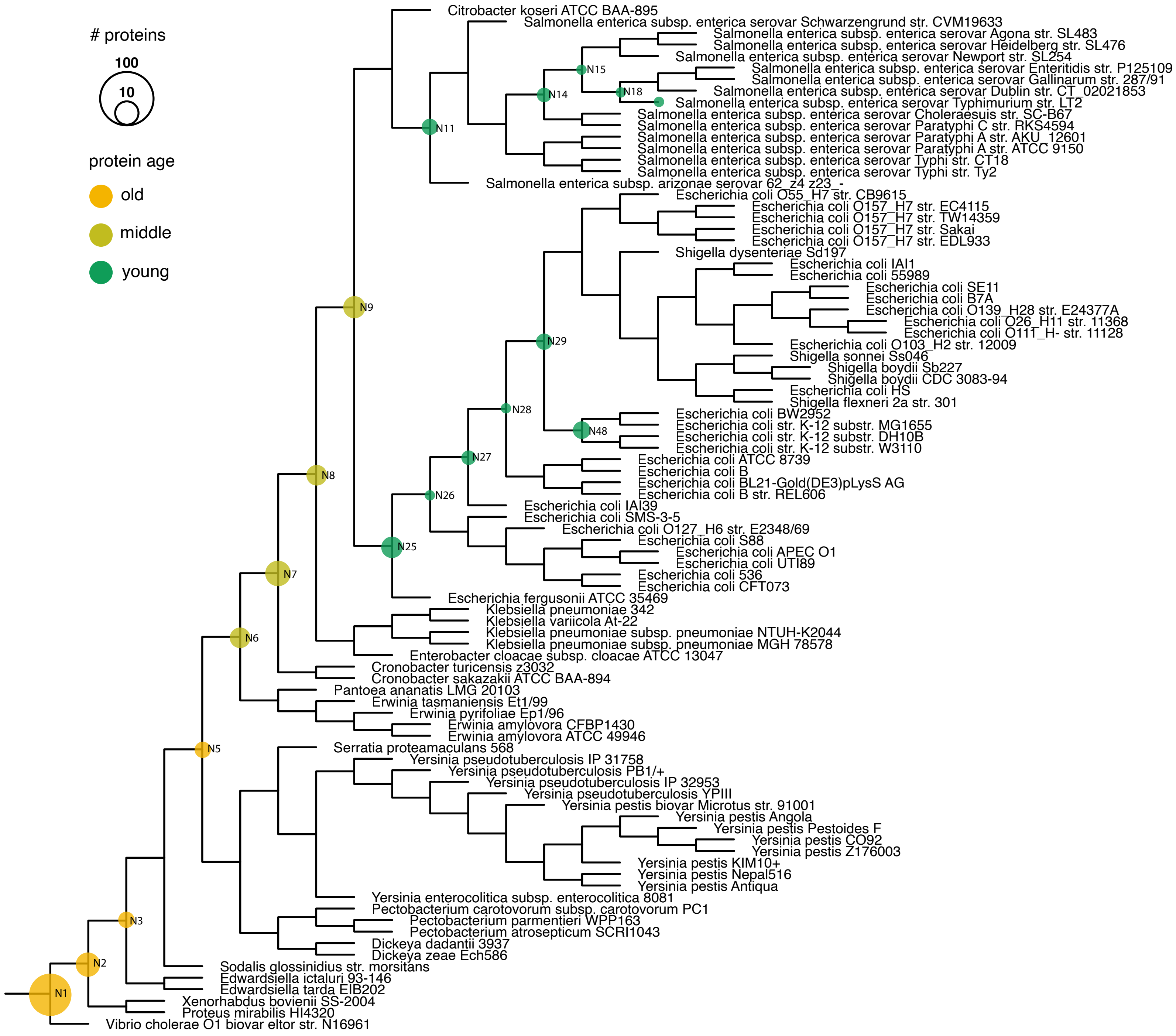

### proteins
100
10
protein age
old
middle
young
Citrobacter koseri ATCC BAA-895
Salmonella enterica subsp. enterica serovar Schwarzengrund str. CVM19633
Salmonella enterica subsp. enterica serovar Agona str. SL483
Salmonella enterica subsp. enterica serovar Heidelberg str. SL476
Salmonella enterica subsp. enterica serovar Newport str. SL254
Salmonella enterica subsp. enterica serovar Enteritidis str. P125109
Salmonella enterica subsp. enterica serovar Gallinarum str. 287/91
Salmonella enterica subsp. enterica serovar Dublin str. CT_02021853
Salmonella enterica subsp. enterica serovar Typhimurium str. LT2
Salmonella enterica subsp. enterica serovar Choleraesuis str. SC-B67
Salmonella enterica subsp. enterica serovar Paratyphi C str. RKS4594
Salmonella enterica subsp. enterica serovar Paratyphi A str. AKU_12601
Salmonella enterica subsp. enterica serovar Paratyphi A str. ATCC 9150
Salmonella enterica subsp. enterica serovar Typhi str. CT18
Salmonella enterica subsp. enterica serovar Typhi str. Ty2
Salmonella enterica subsp. arizonae serovar 62_z4 z23_-
Escherichia coli O55_H7 str. CB9615
Escherichia coli O157_H7 str. EC4115
Escherichia coli O157_H7 str. TW14359
Escherichia coli O157_H7 str. Sakai
Escherichia coli O157_H7 str. EDL933
Shigella dysenteriae Sd197
Escherichia coli IAI1
Escherichia coli 55989
Escherichia coli SE11
Escherichia coli B7A
Escherichia coli O139_H28 str. E24377A
Escherichia coli O26_H11 str. 11368
Escherichia coli O111_H- str. 11128
Escherichia coli O103_H2 str. 12009
Shigella sonnei Ss046
Shigella boydii Sb227
Shigella boydii CDC 3083-94
Escherichia coli HS
Shigella flexneri 2a str. 301
Escherichia coli BW2952
Escherichia coli str. K-12 substr. MG1655
Escherichia coli str. K-12 substr. DH10B
Escherichia coli str. K-12 substr. W3110
Escherichia coli ATCC 8739
Escherichia coli B
Escherichia coli BL21-Gold(DE3)pLysS AG
Escherichia coli B str. REL606
Escherichia coli IAI39
Escherichia coli SMS-3-5
Escherichia coli O127_H6 str. E2348/69
Escherichia coli S88
Escherichia coli APEC O1
Escherichia coli UTI89
Escherichia coli 536
Escherichia coli CFT073
Escherichia fergusonii ATCC 35469
Klebsiella pneumoniae 342
Klebsiella variicola At-22
Klebsiella pneumoniae subsp. pneumoniae NTUH-K2044
Klebsiella pneumoniae subsp. pneumoniae MGH 78578
Enterobacter cloacae subsp. cloacae ATCC 13047
Cronobacter turicensis z3032
Cronobacter sakazakii ATCC BAA-894
Pantoea ananatis LMG 20103
Erwinia tasmaniensis Et1/99
Erwinia pyrifoliae Ep1/96
Erwinia amylovora CFBP1430
Erwinia amylovora ATCC 49946
Serratia proteamaculans 568
Yersinia pseudotuberculosis IP 31758
Yersinia pseudotuberculosis PB1/+
Yersinia pseudotuberculosis IP 32953
Yersinia pseudotuberculosis YPIII
Yersinia pestis biovar Microtus str. 91001
Yersinia pestis Angola
Yersinia pestis Pestoides F
Yersinia pestis CO92
Yersinia pestis Z176003
Yersinia pestis KIM10+
Yersinia pestis Nepal516
Yersinia pestis Antiqua
Yersinia enterocolitica subsp. enterocolitica 8081
Pectobacterium carotovorum subsp. carotovorum PC1
Pectobacterium parmentieri WPP163
Pectobacterium atrosepticum SCRI1043
Dickeya dadantii 3937
Dickeya zeae Ech586
Sodalis glossinidius str. morsitans
Edwardsiella ictaluri 93-146
Edwardsiella tarda EIB202
Xenorhabdus bovienii SS-2004
Proteus mirabilis HI4320
Vibrio cholerae O1 biovar eltor str. N16961
N1
N2
N3
N5
N6
N7
N8
N9
N11
N14
N15
N18
N25
N26
N27
N28
N29
N48

**Figure S2**

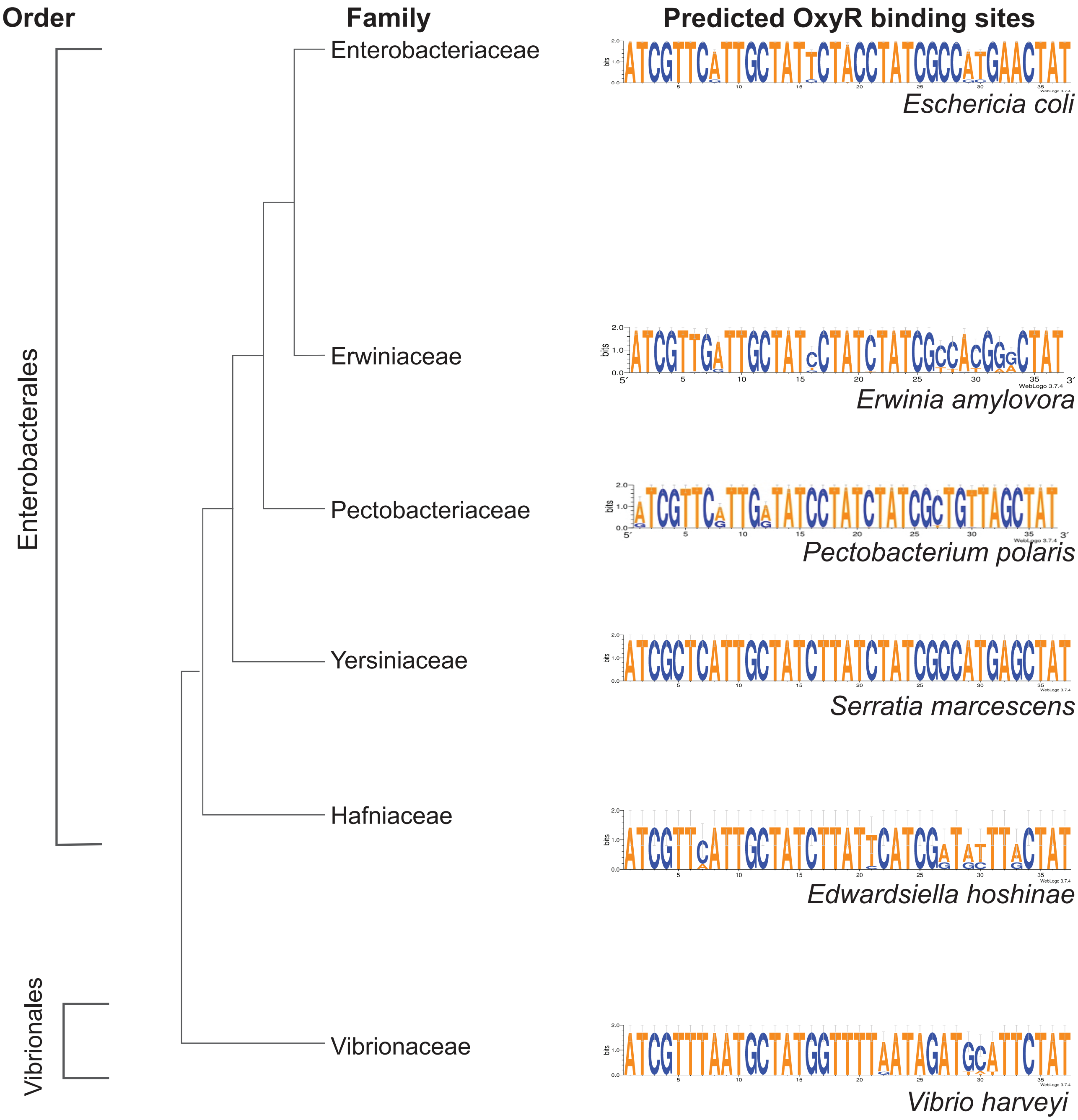

Order
Family
Predicted OxyR binding sites
Enterobacterales
Vibrionales
Enterobacteriaceae
Erwiniaceae
Pectobacteriaceae
Yersiniaceae
Hafniaceae
Vibrionaceae
Eschericia coli
Erwinia amylovora
Pectobacterium polaris
Serratia marcescens
Edwardsiella hoshinae
Vibrio harveyi

#### Figure S3

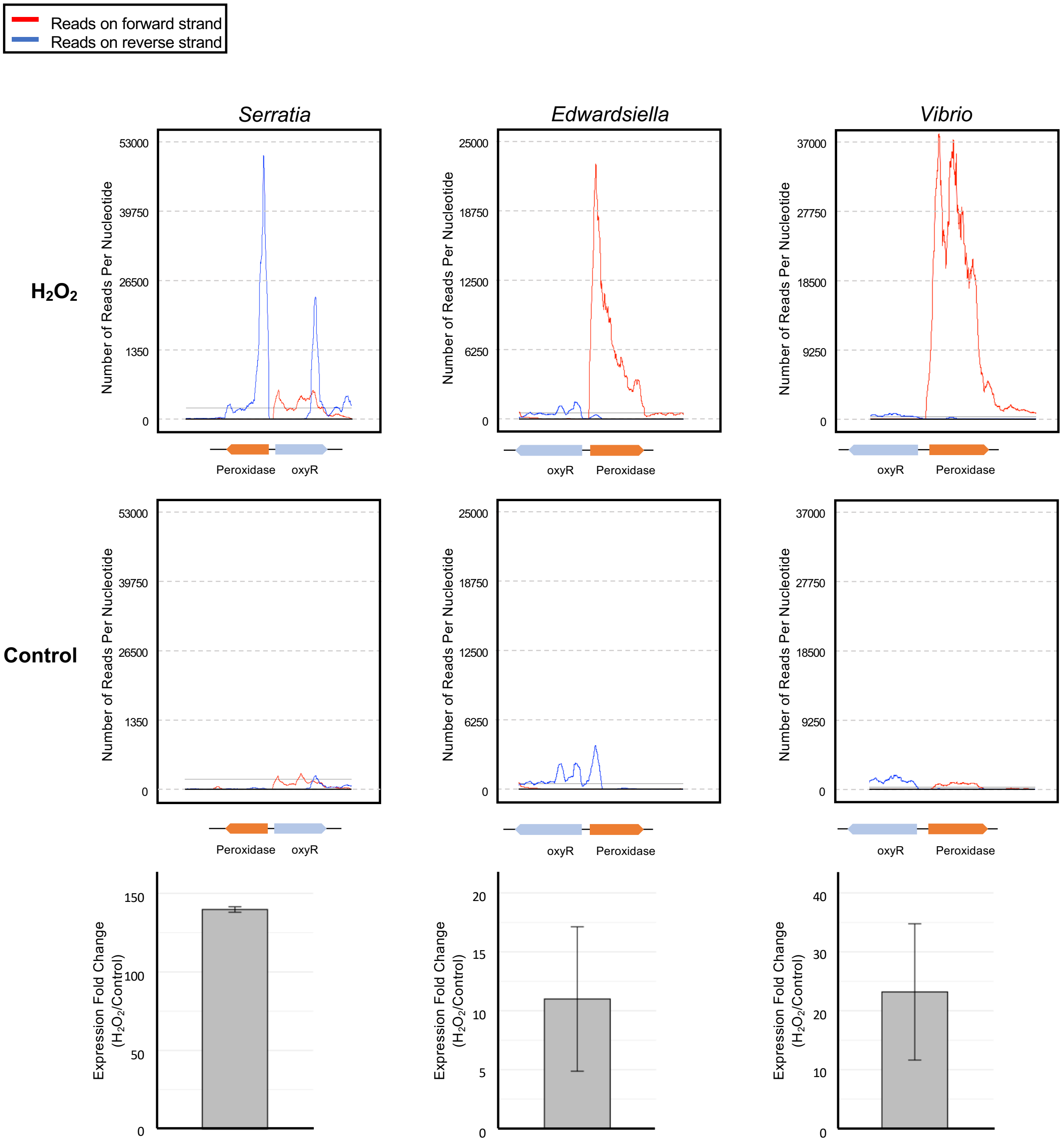

Reads on forward strand
Reads on reverse strand
Serratia
Edwardsiella
Vibrio
$H_2O_2$
Control
Number of Reads Per Nucleotide
53000
39750
26500
1350
0
25000
18750
12500
6250
37000
27750
18500
9250
Peroxidase
oxyR
Expression Fold Change ($H_2O_2$/Control)
150
100
50
20
15
10
5
40
30

**Figure S4**

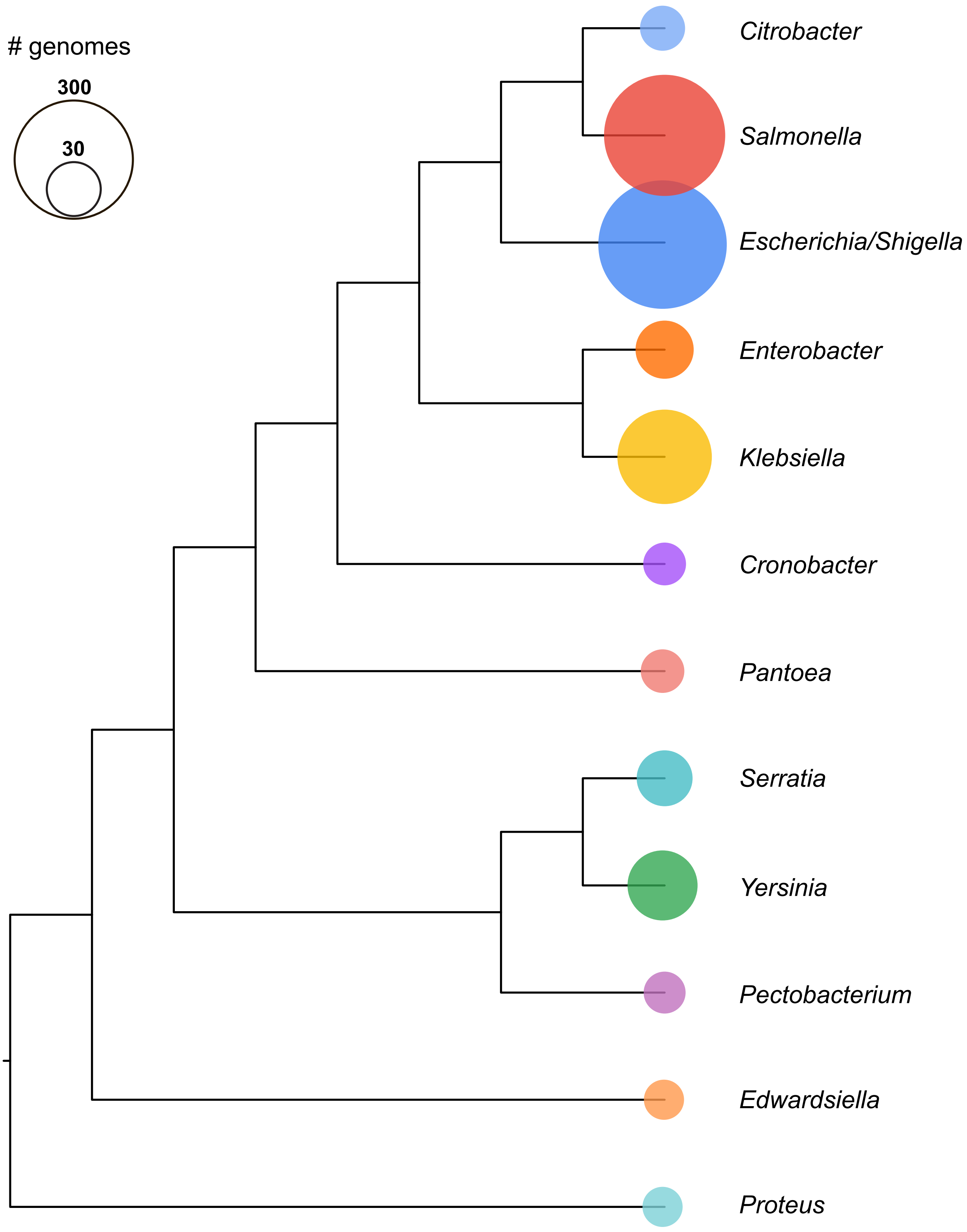

### genomes
300
30
Citrobacter
Salmonella
Escherichia/Shigella
Enterobacter
Klebsiella
Cronobacter
Pantoea
Serratia
Yersinia
Pectobacterium
Edwardsiella
Proteus
